# Rewards interact with explicit knowledge to enhance skilled motor performance

**DOI:** 10.1101/745851

**Authors:** Sean P. Anderson, Tyler J. Adkins, Bradley S. Gary, Taraz G. Lee

## Abstract

From typing on a keyboard to playing the piano, many everyday skills require the ability to quickly and accurately perform sequential movements. It is well-known that the availability of rewards leads to increases in motivational vigor whereby people enhance both the speed and force of their movements. However, in the context of motor skills, it is unclear whether rewards also lead to more effective motor planning and action selection. Here, we trained human participants to perform four separate sequences in a skilled motor sequencing task. Two of these sequences were trained explicitly and performed with pre-cues that allow for the planning of movements, while the other two were trained implicitly. Immediately following the introduction of performance-contingent monetary incentives, participants improved their performance on all sequences consistent with enhancements in motivational vigor. However, there was a much larger performance boost for explicitly trained sequences. We replicated these results in a second, pre-registered experiment with an independent sample. We conclude from these experiments that rewards enhance both the planning of movements as well as motivational vigor.

## 1. Introduction

Rewards and incentives drive people to work more vigorously and to expend more energy to accomplish their goals. This increase in *motivational vigor* has been shown to improve performance of simple motor movements (e.g., reaches, saccades, etc.) by shifting the speed-accuracy trade-off function, resulting in movements that are both faster and more accurate (Manohar et al., 2015; Summerside, Shadmehr, & Ahmed, 2018; Takikawa, Kawagoe, Itoh, Nakahara, & Hikosaka, 2002). Motor skills, and specifically motor sequencing skills, involve the precise execution of a chain of simple motor movements at specific times. However, they also require recognition of the current context and the rapid selection of the correct action given that context. As a sequencing skill develops, these processes can be pre-planned, allowing for improved performance (Ariani & Diedrichsen, 2019; Diedrichsen & Kornysheva, 2015). While reward availability and increased motivation also improve performance of more complex motor skills (Mosberger, De Clauser, Kasper, & Schwab, 2016; Wachter, Lungu, Liu, Willingham, & Ashe, 2009), it remains unclear which stages of the motor planning hierarchy from stimulus to movement (action selection, action execution, etc.) are sensitive to these motivational effects.

Motor sequence learning tasks such as the serial reaction time task (SRTT) and the discrete sequence production (DSP) task are widely used as a model of motor skill acquisition (see Krakauer, Hadjiosif, Xu, Wong, & Haith, 2019 for a comprehensive review). Although skill acquisition in sequence learning tasks is often thought to be largely implicit (Nissen & Bullemer, 1987; Reber & Squire, 1994), these skills can be taught at least partially via explicit instruction (Grafton, Hazeltine, & Ivry, 1995; D. B. Willingham & Goedert-Eschmann, 1999). For example, providing the sequence or alerting the learner that a to-be-learned sequence is present can result in faster acquisition, fewer errors produced, and more rapid execution (Lee, Acuña, Kording, & Grafton, 2019; Schendan, Searl, Melrose, & Stern, 2003; Daniel B Willingham, Salidis, & Gabrieli, 2002). Traditionally, it has been thought that explicit knowledge of sequence order enables advanced motor planning of the known sequence elements, essentially enhancing the action selection process (Diedrichsen & Kornysheva, 2015; Hikosaka, Nakamura, Sakai, & Nakahara, 2002). Explicit training is also associated with cognitive control, a collection of cognitive processes that promote goal-directed behavior. For example, larger working memory capacity increases the rate of explicit skill learning (Anguera, Reuter-Lorenz, Willingham, & Seidler, 2010) and explicit training coincides with increases in activity in brain regions such as the dorsolateral prefrontal cortex that are important for working memory and other cognitive control processes (Grafton et al., 1995; Schendan et al., 2003; Daniel B Willingham et al., 2002). Performance-contingent rewards have been shown to enhance cognitive control processes. In particular, rewards have been associated with improvements in proactive control, which requires the advanced maintenance of goal information (Botvinick & Braver, 2014; Etzel, Cole, Zacks, Kay, & Braver, 2015; Jimura, Locke, & Braver, 2010; Padmala & Pessoa, 2011). Thus, reward-based enhancement of motor skills could be due in part to enhancements in cognitive control and prospective planning in addition to improvements in simple motor execution.

However, some recent evidence suggests that explicit sequence knowledge does not affect motor planning, but instead results in an overall increase in motivation (Wong et al., 2015). This increase in motivation in turn globally enhances movement vigor. In this study, participants were faster at executing known sequences than random sequences, but they showed similar performance improvements in executing movements for random elements embedded within these known sequences. Since the participants could not plan the random elements, their improvements could not be due to benefits in movement planning. This result suggests that explicit knowledge improves skilled performance by increasing the motivational vigor of all movements during execution rather than by enabling the advanced planning of specific goal-oriented movements.

Here, we sought to determine whether motivational enhancements of skilled performance result from an improvement in motor execution, an improvement in motor planning, or both. Across two experiments, human participants were trained on four separate motor sequences in a discrete sequence production (DSP) task. Two of the sequences were implicitly trained, while the other two were trained with explicit instructions and pre-movement cues that could allow participants to plan movement sequences in advance. Immediately following training, participants performed these learned skills for cash rewards to assess how increased motivation impacts skilled motor performance. We hypothesized that if monetary incentives improve skilled performance by simple increases in motivational vigor for all movements, we would observe similar levels of improvement for both implicitly and explicitly trained skills. If, additionally, explicit knowledge itself acts as a reward cue, this would increase motivational vigor but not aid motor planning, as argued by Wong et al. (2015). In this case one might expect that the introduction of monetary incentives would disproportionally benefit implicitly trained skills relative to explicitly trained skills. That is, implicit skills would have more room to improve upon the introduction of external motivation via incentives as explicit skills already received a motivational boost due to explicit knowledge. This assumes there is a point at which improvements in motivational vigor saturate, and that explicit knowledge and reward improve performance through similar mechanisms (i.e. enhanced motivational vigor). In contrast to the above, we expected that if explicit training leads to increases in skill knowledge that facilitates motor planning, monetary incentives would lead to larger improvements in explicit skills relative to implicit skills. To validate our results following Experiment 1, we pre-registered our hypotheses and analyses and performed a direct replication of this initial experiment in a separate sample of individuals (Experiment 2).

## 2. EXPERIMENT 1

### 2.1. Experiment 1 Materials and Methods

#### 2.1.1. Participants

61 Participants participated in Experiment 1 (48 Females, 12 Males, 1 unreported, all right-hand dominant, mean age = 20.8). 4 Participants were excluded from the entire experiment because they used both hands. 1 additional participant left to use the bathroom in the middle of the experiment; their data was excluded from the analyses. Data was used from 56 participants in total. However, an additional subject was excluded from Generation and Recognition analyses (see Explicit Knowledge Tests below) because they left the experiment before the memory tests could be completed leaving 55 participants in those analyses. All participants were paid $10/hr + performance bonuses for their participation. All participants provided written, informed consent and all research protocols were improved by the Health Sciences and Behavioral Sciences Institutional Review Board at the University of Michigan.

#### 2.1.2. Discrete Sequence Production Task

The task consisted of a modification of the Discrete Sequence Production task (Verwey, Lammens, & van Honk, 2002). Participants sat centered in front of a computer screen with a standard QWERTY keyboard and were instructed to use their non-dominant left hand at all times. During a single trial, a colored square (see below) was displayed in the center of the screen for 1 s to notify participants of the identity of the upcoming sequence. Afterwards, four grey and square placeholders were displayed corresponding to four adjacent keys (A, S, D, F) for another 1 s. One of the placeholder boxes turned white until the corresponding key was pressed, after which it would be extinguished and another placeholder box would turn white. This continued for a total of 8 items per trial, making up a sequence. Participants were instructed to type these sequences as quickly and as accurately as possible and that they would be eligible to receive cash bonuses for their performance in a separate phase of the experiment. If an incorrect key was pressed, the corresponding placeholder box would turn red for one second and the experiment advanced to the next trial. There was also a time deadline for each trial: set to 8 s during Training and calibrated to the individual during the Reward phase (see below). If the participant did not successfully type the entire 8-item sequence before the time deadline a message saying “Too Slow” was displayed for one second and the trial was aborted.

#### 2.1.3. Training phase

Each participant was exposed to four separate 8-item sequences picked from a corpus of possible sequences. Each sequence did not include trills (e.g. 1-2-1-2) nor runs (e.g. 1-2-3-4) and began with a different key. Two of these sequences were presented with a unique color cue appearing at the start of the trial to facilitate explicit training. Participants were told that they were to learn to associate the color cues with the sequences and that this would aid in training. A gray cue appeared before each of the remaining two sequences. Participants were told that the gray cue signified that the upcoming trial would be randomly generated. These sequences act as a control to allow us to compare to the cued sequences to measure general skill improvement. To ensure that these sequences were learned implicitly, on a subset of trials participants were exposed to a gray cue followed by a pseudo-randomly generated sequence to prevent participants from noticing the repeating patterns (see Figure 1a).

**Figure 1.**
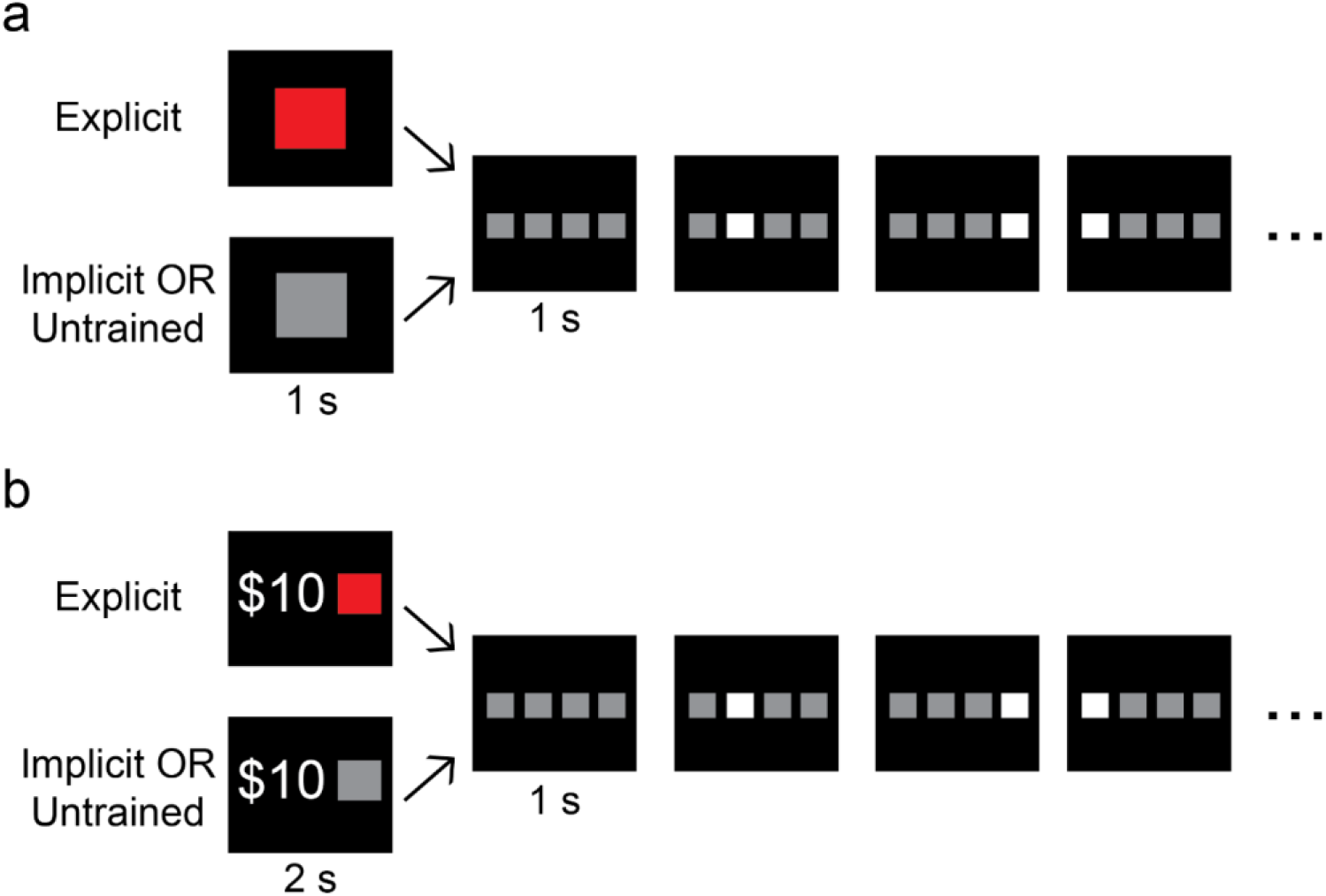
Discrete Sequence Production Task. a) Participants trained to perform four different 8-item sequences (two explicit, two implicit). Prior to execution of explicitly trained sequences, a colored square was presented to cue the participants of the identity of the upcoming sequence to facilitate awareness and to allow for movement pre-planning. For both implicitly trained sequences, a gray square was presented. Participants were told that sequences following a gray cue would be randomly generated. b) Immediately following training, participants performed the same task for cash bonuses. The reward value ($5, $10, or $20) for each trial appeared simultaneously with the sequence cue during this phase of the experiment.

Participants completed a total of 400 trials during training. Within the two Explicit conditions, one sequence was trained extensively (Deep), appearing in 120 trials during the Training phase. The other Explicit sequence appeared in only 40 trials during Training (Shallow). Similarly, one of the Implicit sequences appeared on 120 trials (Deep) and the other just 40 trials (Shallow). The pseudo-randomly generated sequences (Untrained) appeared on 80 trials (20% of the total number of trials). All together there were five conditions: Explicit/Deep (E/D), Explicit/Shallow (E/S), Implicit/Deep (I/D), Implicit/Shallow (I/S), and Untrained. The actual sequences used for each of these conditions was counterbalanced across participants such that no observed effects between conditions could be due to the precise sequences performed. Each participant completed 8 blocks of 50 trials each during training. The order of each sequence presentation was randomized within each block. Prior to training, participants were told that there would be a “test” phase following training to assess how much they had learned.

#### 2.1.4. Reward phase

After the Training phase, participants were informed that they would now complete the task for the chance to win cash bonuses. During the cue presentation (2 seconds) at the outset of each trial, an incentive value for the trial was displayed: $5, $10, or $20 (Figure 1b). Participants were instructed that at the end of the experiment, a trial would be selected at random, and if they successfully completed that trial (pressing all 8 items within the time limit) they would receive the associated reward for that trial. This encouraged participants to evaluate each incentive independently of the other trials.

Participants were also instructed that we would impose a stricter time limit during the Reward phase to ensure that they didn’t slow down to try to earn rewards. Participants completed 9 blocks of 27 trials each. The sequences (E/D, E/S, I/D, I/S) were presented on 48 trials each, 16 for each incentive value. Untrained sequences appeared in 51 trials. To protect against carryover effects, incentive values were presented in an m-sequence (Buračas & Boynton, 2002).

In order to equalize trial difficulty across participants and sequence conditions, time limits for each sequence were calibrated individually based on participants’ performance at the end of the Training phase. For each participant, the time limit for each sequence was calculated by taking the 75^th^ percentile of their movement times during their last eight accurate trials in Training for that sequence. Each subject had a different time limit for each of the sequence conditions. This enabled us to use accuracy as a dependent measure and compare it across participants and conditions. Additionally, this reduced the possibility of floor and ceiling effects, and minimized speed/accuracy trade-offs during performance. That is, the time limits prevented participants from purposely slowing their execution time in order to be more accurate and increase their chances of success.

#### 2.1.5. Explicit Knowledge tests

After the Reward phase, participants completed two tests to assess their explicit knowledge of the sequences. In a free recall sequence generation test, participants were informed that not all grey-cued sequences were random, and there were in fact two sequences and a random condition during grey cued trials. Participants were then asked to type out each sequence from memory. For the explicitly cued sequences, participants attempted to type the sequence after the matching color cue was displayed. Since all sequences began with a unique key, participants were given the first item of the sequence for the implicitly trained sequences and asked to type the rest. Performance in the generation test was assessed by computing the Damerau-Levenshtein edit-distance (Damerau, 1964; Levenshtein, 1966) between the typed sequence and the correct sequence. This value is the minimum number of translations (insertions, deletions, transpositions, or substitutions) needed to transform one sequence into the other. Because participants were given the first key for each of the implicit sequences, this measure likely slightly over-estimates recall for these sequences.

Immediately following the free recall test, participants completed a recognition test. Participants were prompted to complete a total of 16 separate sequences and asked to rate on a 7-point Likert scale how likely the sequences were to be new or old. 4 of the sequences were the trained ones from the experiment and 12 were randomly generated. Explicit knowledge was assessed by calculating the difference between the mean rating for random sequences to that for each trained sequence. Larger numbers indicate a greater ability to distinguish untrained and trained sequences.

#### 2.1.6. Data Analysis

During each trial, response times for each item in the sequence were recorded. Movement Time (MT) was calculated as the duration between the first and last button presses. The trial was considered correct if all 8 items were pressed correctly before the time limit. Accuracy was defined as the proportion of correct trials within the condition(s) in question. Response Time (RT) was calculated as the duration between the presentation of the first item (when the placeholder box turned white) and the first button press. Inter-Key Interval (IKI) for each keypress was calculated as the duration between the previous keypress and the current keypress (i.e. IKI for item 3 was the time between item 2 being pressed and item 3 being pressed). As responses were recorded on a standard keyboard, we did not obtain measures of force. All analyses involving MT, RT, and IKI were performed on accurate trials only. Additionally, to assess the immediate effect of introducing incentives, we computed the percentage change in performance (MT, RT, and each IKI) between the last trials of the Training phase and the first trials of the Reward phase. Specifically, for each subject and each sequence condition, we computed the percent change between the mean performance on the last 10 accurate trials of Training (Last10_*T*_) and the mean performance on the first 10 accurate trials of Reward (First10_*R*_):

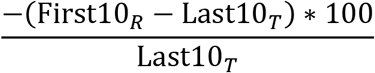

Importantly, we chose to analyze percentage improvement instead of the raw absolute change of MT and RT in order to more fairly assess the relative enhancements in performance for each condition. For instance, if an Olympic track athlete improves an already fast lap time by a few seconds, their improvement likely reflects a more impressive level of improvement when compared to a novice (i.e. slow) runner improving their time by that same amount. Essentially, the percentage improvement measure has the advantage of normalizing performance enhancements against individual levels of performance at the end of the Training phase, whereas absolute change measures ignore previous performance.

The choice of exactly 10 trials was somewhat arbitrary, but was selected to provide a meaningful measure of performance at the end of training. As there were only 5 trials per block for each of the shallow sequences, including more than 10 correct trials from the end of Training would have had a high likelihood of including trials that occurred toward the middle of the Training phase of the experiment. This would likely inflate the estimate of reward-related enhancement for shallow trials with contamination from sequence-independent improvement in movement time across the last several blocks of training. However, we performed control analyses to ensure our results were robust to the number of trials selected (see Supplemental Material).

To examine the effect of explicit cues and training depth on sequence learning during training, we entered training block number, cue type (explicit vs. implicit) and training depth (Deep vs. Shallow) as within-subject factors in separate repeated-measures ANOVAs for each dependent variable of interest (RT on correct trials, MT on correct trials, accuracy).

To examine the effect of explicit cues and training depth on percentage improvement (between Last10_*T*_ and First10_*R*_), we entered cue type and training depth as within-subject factors in separate repeated-measures ANOVAs on percentage improvement in MT and in RT. To examine the effect of explicit cues on percentage improvement in IKI, we averaged percentage improvement across the two explicit sequences and the two implicit sequences for each button press. We then performed dependent t-tests at each IKI (2-8), correcting for multiple comparisons via the Holm-Bonferroni method.

To examine the effect of explicit cues, training depth, and incentive value on performance during Reward, we entered cue type, training depth, and incentive value ($5, $10, or $20) as within-subject factors in separate repeated-measures ANOVAs for each dependent variable of interest.

To examine the effect of explicit cues and training depth on knowledge of the sequences, we entered cue type and training depth as within-subject factors in separate repeated-measures ANOVAs on generation test performance and recognition test performance. In all ANOVAs, violations of sphericity were corrected using the Greenhouse-Geiser method.

### 2.2. Experiment 1 Results

#### 2.2.1. Explicit cues and increased practice led to better learning and performance in Training

We first sought to verify the effectiveness of explicit cues in enhancing motor sequence learning. As expected, participants’ motor skill improved throughout training, as they greatly reduced their total movement time (MT) for each sequence (Main effect of Block: F_3.96, 209.71_ = 163.390, p < 0.001, η_p_^2^ = 0.755; Figure 2a) and also were better able to complete the 8-item sequences without execution errors (Main effect of Block on accuracy rate: F_5.03, 276.73_ = 2.32, p = 0.043, η_p_^2^ = 0.04). MT was faster overall for explicitly cued sequences relative to implicitly cued sequences (main effect of explicit cues: F_1,53_ = 30.982, p < 0.001, η_p_^2^ = 0.369). Explicit cues were also associated with a faster rate of improvement in MT as training progressed (cue x block interaction: F_7,371_ = 27.434, p < 0.001, η_p_^2^ = 0.341). Similarly, explicit cues allowed for faster response times (RT) to the first item of each sequence (F_1,55_ = 67.806, p < 0.001, η_p_^2^ = 0.552) and this advantage for explicitly cued sequences grew larger over the course of training (cue x block interaction: F_2.927, 160.994_ = 8.961, p < 0.001, η_p_^2^ = 0.140). While there was not a main effect of explicit cueing on overall accuracy rate (F_1,55_ = 0.177, p = 0.676, η_p_^2^ = 0.003), we did find a cue x block interaction (F_7, 385_ = 2.935, p = 0.005, η_p_^2^ = 0.051) driven by the fact that accuracy in explicit sequences improved at a slightly faster rate toward the end of training (see Figure S1). These results point to tangible enhancements in learning and performance due to the explicit sequence cues rather than a simple shift along the speed-accuracy tradeoff curve (i.e. faster speed, but reduced accuracy on explicit sequences). Explicit cues during training conferred performance benefits to both learning and performance consistent with prior work.

**Figure 2.**
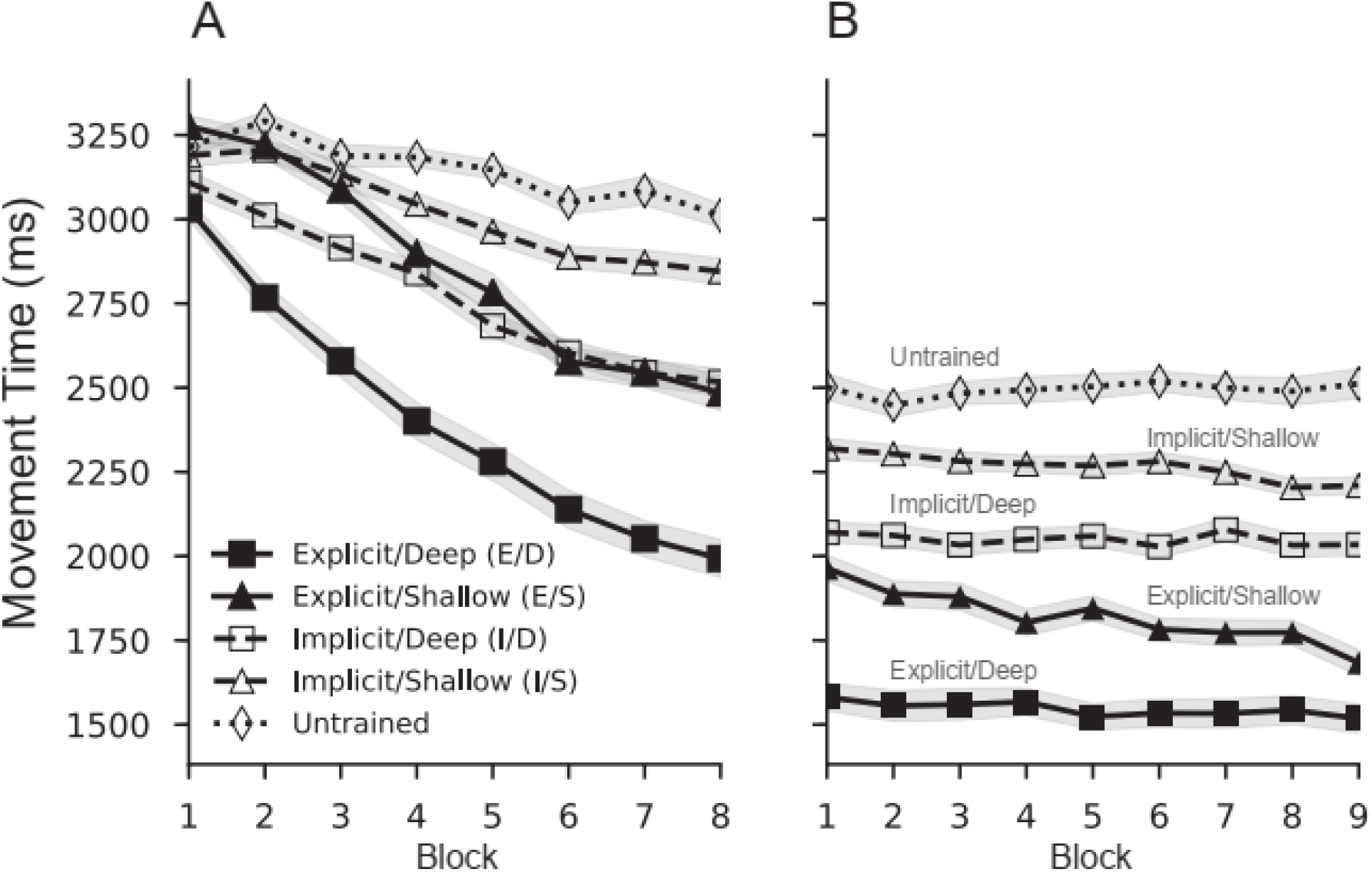
Mean movement time (MT) for correctly executed sequences. MT was measured as the total time between the first and last item in each sequence. a) Explicit sequence cues and increased training were both associated with faster rates of learning. b) Following the introduction of performance-contingent monetary incentives, movement times immediately decreased for all sequences. Shaded areas indicate SEM^norm^. (E/D - Deeply trained, Explicit; I/D - Deeply trained, Implicit, E/S - Shallowly trained, Explicit, I/S - Shallowly trained, Implicit).

We also sought to validate the effect of practice (depth of training) on performance. MT was faster for the deeply trained sequences (main effect of depth: F_1,53_ = 214.23, p < 0.001, η_p_^2^ = 0.802) and the size of this advantage grew over the course of training (Block x Depth interaction: F_4.802_ = 9.570, p < 0.001, η_p_^2^ = 0.153). Initial RT was also faster for deeply trained sequences relative to shallowly trained sequences (F_1,55_ = 6.609, p = 0.013, η_p_^2^ = 0.107). Accuracy was also higher for deeply trained sequences (F_1,55_ = 66.941, p < 0.001, η_p_^2^ = 0.549). Additionally, we observed a knowledge type by training depth interaction in MT (F_1,53_ = 21.846, p < 0.001, η_p_^2^ = 0.292). The benefit of training depth on MT was more pronounced for explicitly cued sequences than for implicitly trained sequences. This interaction does not appear to be driven by a speed-accuracy tradeoff (faster MT, but reduced accuracy) as we did not observe a knowledge type by training depth interaction in accuracy (F_1,55_ = 0.065, p = 0.800 η_p_^2^ = 0.001). In aggregate, and perhaps unsurprisingly, more practice on a sequence corresponded with better performance.

#### 2.2.2. Monetary incentives lead to larger immediate performance enhancement for skills trained explicitly rather than implicitly

We were primarily interested in whether explicit knowledge moderates the effect of motivation on motor performance through improved motor planning. To this end, we examined the immediate effect of the introduction of performance-contingent monetary incentives following the Training phase of the experiment. Any change in performance between the two phases of the experiment is unlikely due to learning given the limited number of additional trials; rather, it should reflect immediate enhancements in motor vigor and/or motor planning due to increased motivation. Here, we use percent change in performance as an index of reward-related enhancement as skill level differed markedly between conditions (see Methods). In response to this increase in extrinsic motivation, participants were able to immediately enhance the speed of their execution for all trained sequences and Untrained sequences (see Figure 2b and Figure 3; p< 0.001 in one-sample t-tests for all sequences). The fact that there was a large decrease in MT on Untrained sequences (19% on average) suggests that monetary incentives had a global effect on motivational vigor independent from any sequence-specific skill knowledge. The size of this boost in performance was virtually identical for Untrained sequences and the two implicitly trained sequences (no main effect of sequence identity, explicit sequences excluded: F_1.720, 92.904_ = 1.254, p = 0.286, η_p_^2^ = 0.023). However, MT improvement for explicitly cued sequences was significantly greater than that for both implicitly trained sequences (main effect of cue type: F_1,55_ = 33.485, p < 0.001, η_p_^2^ = 0.378) and Untrained sequences (E/D vs. Untrained: t_55_ = 3.50, p = 0.002; E/S vs. Untrained: t_55_ = 5.88, p < 0.001). None of these findings were driven by the number of trials used for this analysis; similar results were obtained across all comparisons when running the analysis using 5 or 15 trials from each phase of the experiment (all p-values less than 0.05). These results suggest that monetary incentives produced skill enhancements for explicitly trained sequences above and beyond simple increases in overall motor vigor.

**Figure 3.**
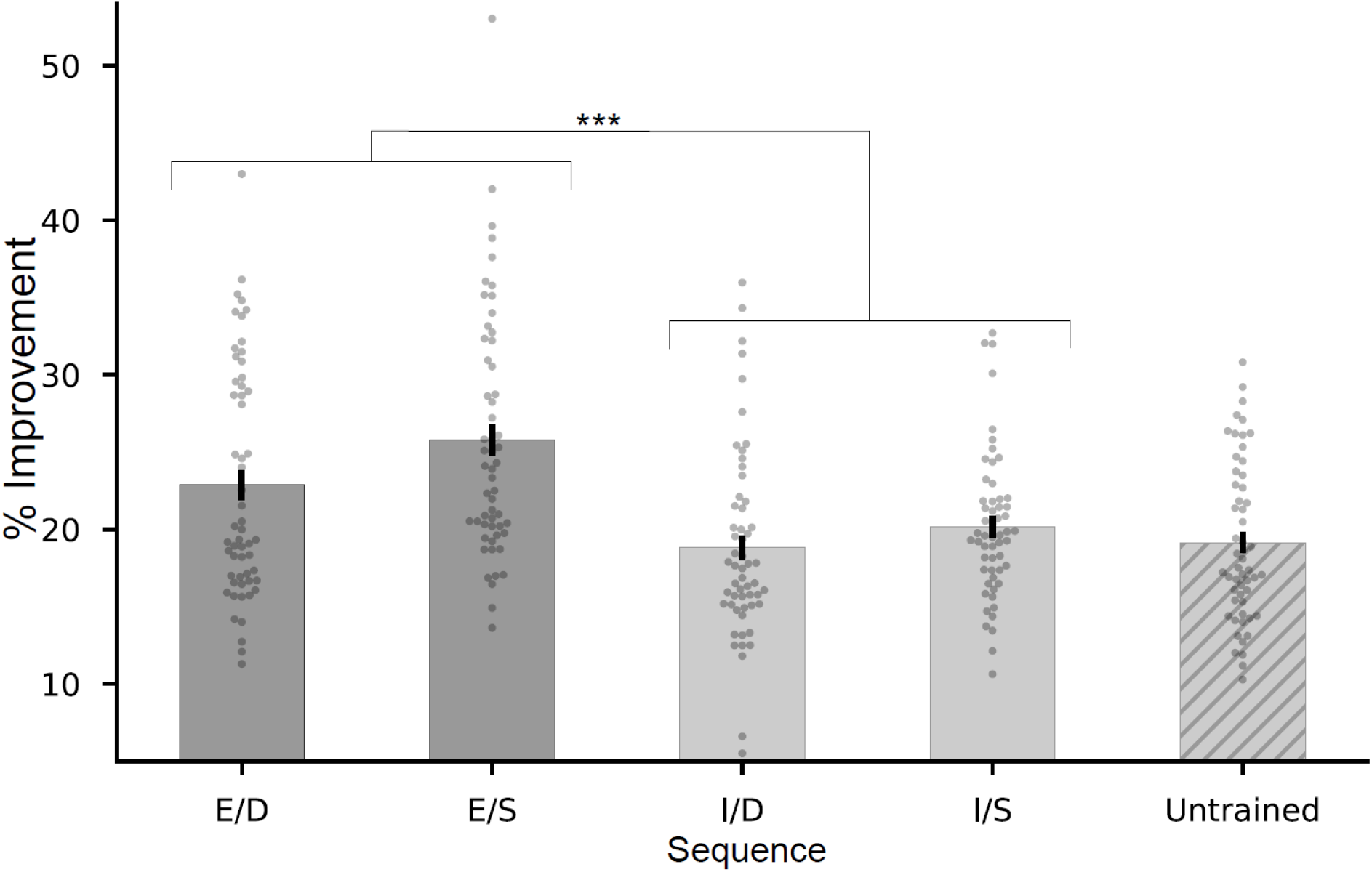
Incentive-based performance improvements. Immediately following the introduction of performance-contingent monetary incentives, participants improved their performance of all sequences. While the size of this performance improvement was similar for both implicit sequences and untrained sequences, explicitly trained skills improved to a much larger extent. Gray dots indicate individual subjects. Error bars indicate SEM^norm^. (E/D - Deeply trained, Explicit; I/D - Deeply trained, Implicit, E/S - Shallowly trained, Explicit, I/S - Shallowly trained, Implicit; *** = p < 0.001).

These effects cannot simply be explained by the fact that participants are relatively faster at the end of training for the explicitly cued sequences. We did not find a statistically significant relationship between MT at the end of training and the size of the reward-related performance enhancement for any of the sequences following correction for multiple comparisons via the Holm-Bonferroni method (E/D: r_s_ = 0.13, p_uncorrected_ = 0.33, p_corrected_ = 0.52; E/S: r_s_ = 0.22, p_unc_ = 0.10, p_c_ = 0.40; I/D: r_s_ = −0.15, p_unc_ = 0.26, p_c_ = 0.52; I/S: r_s_ = −0.27, p_unc_ = 0.04, p_c_ = 0.20; Untrained: r_s_ = −0.21, p_unc_ = 0.11, p_c_ = 0.40). While we did find evidence that more shallowly trained sequences showed a greater motivational boost overall (main effect of training depth: F_1, 55_ = 8.287, p = 0.006, η_p_^2^ = 0.131) there was no depth x cue type interaction (F_1,55_ = 0.979, p = 0.327, η_p_^2^ = 0.017). Furthermore, participants performed similarly during the last block of training on the I/D sequence and the E/S sequence (t_55_ = 0.483, p = 0.631), yet the immediate reward-based performance improvement was larger for the explicitly trained sequence (E/S) (t_55_ = −6.185, p < 0.001). Additionally, even when measured in raw ms, the immediate shift in MT was larger for E/S than I/D (t_55_ = 5.67 p < 0.001). This rules out the possibility that performance enhancements were due to a simple static shift in execution time across all sequences. Because these sequences were matched in performance prior to the introduction of monetary incentives, this again suggests that increased motivation benefits explicitly trained skills above and beyond a simple increase in motor vigor.

The introduction of monetary incentives also led to faster RT to the first item of each sequence (p < 0.001 in one-sample t-tests for all sequences). However, this reduction in RT was similar across all sequences as we found no significant effects of cue type or training depth (main effect of cue type: F_1,55_ = 0.298, p = 0.587, η_p_^2^ = 0.005; main effect of training depth: F_1,55_ = 0.093, p = 0.761, η_p_^2^ = 0.002; depth x cue interaction: F = 3.459, p = 0.068, η_p_^2^ = 0.059). This result again points to an increase in motivational vigor due to the introduction of monetary incentives. It also suggests that the greater improvement in MT for explicit sequences is due to benefits in performing movements beyond the first item in the sequence.

To more closely examine the possibility that incentive-driven performance enhancements in explicit skills were due to improvements in motor planning, we next examined percent improvement for each inter-key interval (IKI) in explicit and implicit sequences. One might expect greater enhancement for items occurring earlier on in each sequence if cue-based planning were improved by the prospect of rewards. In line with this hypothesis, we observed a significantly larger percentage improvement in explicit sequences relative to implicit sequences for IKIs 2 (between the first keypress and second keypress) and 4 (paired t-test, each p < 0.001, Bonferroni-corrected). We additionally observed a larger performance enhancement for explicit sequences on the final IKI of the sequence (p < 0.001; see Figure 4). No other comparison survived multiple comparisons correction. For the implicitly trained sequences, the incentive-related improvement among all IKIs was similar and not significantly different from the improvement seen for the first item in the sequence (all p > 0.05, corrected). This latter result reinforces the notion that monetary incentives led to a global increase in movement vigor for all items in implicitly trained sequences.

**Figure 4.**
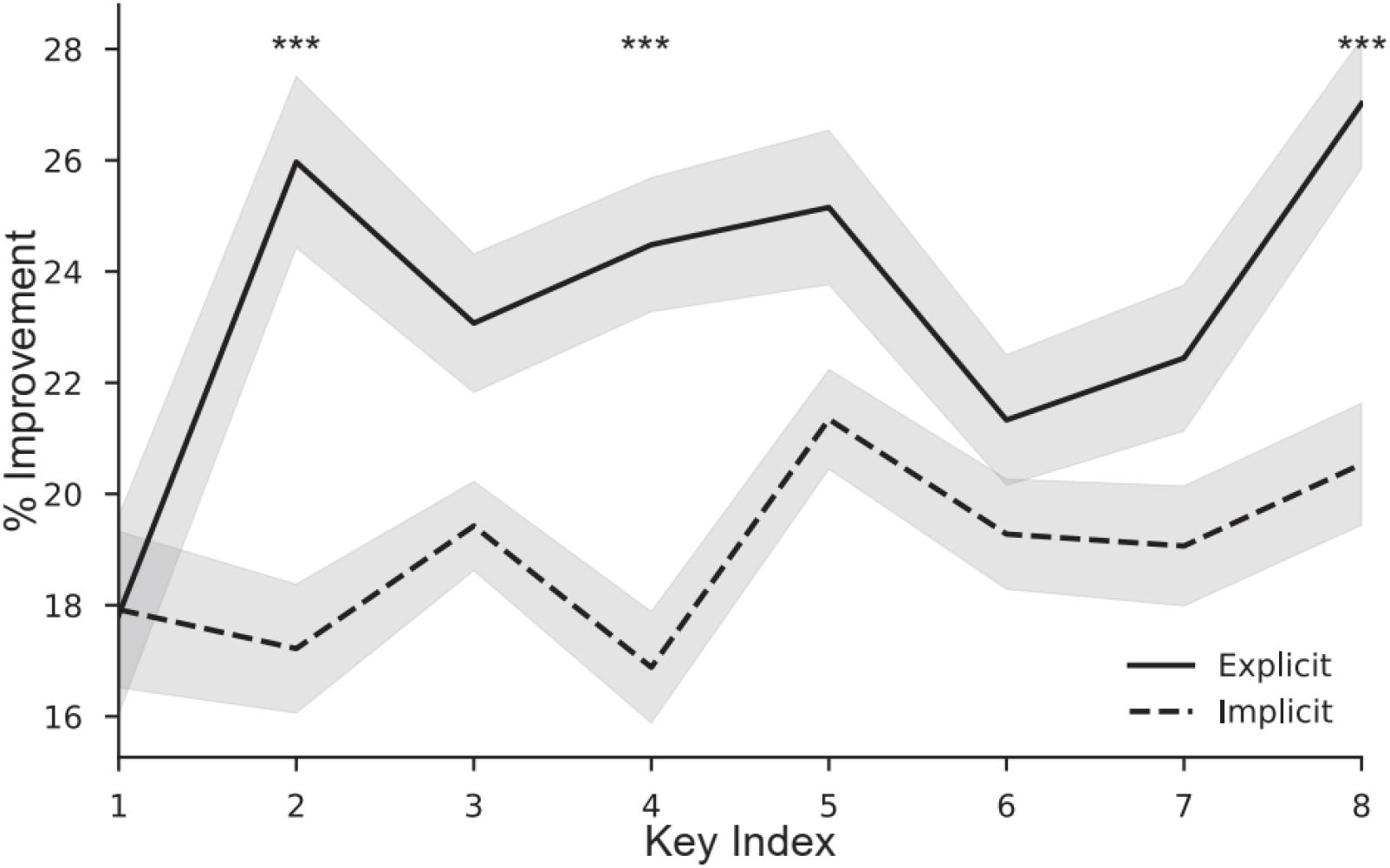
Incentive-based performance improvements for individual items within a sequence. Although all individual movements within a sequence benefitted from the introduction of performance-contingent monetary incentives, items earlier in the sequence (IKI 2 and IKI 4) and at the very end of the sequence (IKI 8) showed significantly larger boosts for explicitly cued sequences. (*** = p < 0.001, Bonferroni-corrected. No other significant differences.)

#### 2.2.3. The benefit of explicit cues and training depth persists with performance-contingent incentives

To verify the performance benefits of explicit knowledge and increased practice remained in contexts with increased motivation, we examined overall performance in the Reward phase of the experiment (Figure 2b). Explicit cues led to faster MT in Reward (F_1,52_ = 77.924, p < 0.001, η_p_^2^ = 0.600). Explicit cues also led to faster RT in Reward (F_1,55_ = 150.125, p < 0.001, η_p_^2^ = 0.732). In other words, the faster MT and RT observed during Training for explicitly cued sequences persisted after the introduction of monetary incentives. MTs were faster in deeply trained sequences during Reward (main effect of training depth: F_1,52_ = 103.403, p < 0.001, η_p_^2^ = 0.665). The same effect was observed in RT (F_1,55_ = 15.997, p < 0.001, η_p_^2^ = 0.225). The overall benefits of explicit cues and depth of training observed during Training remained with the addition of monetary incentives.

#### 2.2.4 Incentive magnitude weakly affects performance accuracy

We next assessed whether performance accuracy was sensitive to incentive magnitude in the Reward phase. Participants were not sensitive to incentive magnitude (main effect of incentive magnitude: F_2,110_ = 0.786, p = 0.458, η_p_^2^ = 0.014). We did observe a relatively small, but significant interaction between reward and training depth on accuracy (F_2,110_ = 4.415, p = 0.014, η_p_^2^ = 0.074). Accuracy for shallowly trained sequences exhibits an inverted-U shape, with relatively greater accuracy for $10 trials (simple main effect of reward for Shallow: F_2,110_ = 3.45, p = 0.035; quadratic contrast: t = 2.58, p = 0.01). Incentive magnitude did not affect accuracy for deeply trained sequences (simple main effect of reward for Deep: F_2,110_ = 1.61, p = 0.21).

We also examined whether training depth or explicit cues resulted in accuracy differences during the Reward phase. During the reward phase, shallowly trained sequences were more accurate overall (main effect of training depth: F_1,55_ = 11.495, p = 0.001, η_p_^2^ = 0.173). Additionally, performance in implicit sequences may have been more accurate than in explicit sequences (marginal effect of cue type: F_1,55_ = 3.781, p = 0.057, η_p_^2^ = 0.064). Given that the time limits were set based on performance at the end of training, these main effects are likely driven by the fact that time limits were somewhat stricter for the more well-learned and explicitly trained sequences.

#### 2.2.5. Greater explicit knowledge for explicitly cued sequences

A primary purpose of the explicit cues was to promote the formation of explicit knowledge of the cued sequences. To assess the effectiveness of this manipulation, we compared accuracy on the free-recall test across the different sequences. Participants were given either the sequence cue (explicit sequences) or the first item of the sequence (implicit sequences) and were asked to key in the entire sequence from memory. As expected, participants were able to recall and execute explicitly cued sequences more accurately than implicitly trained ones (F_1,54_ = 59.259, p < 0.001, η_p_^2^ = 0.523; Figure 5). This is despite the fact that memory for implicit sequences was likely somewhat over-estimated since they were given the first item (see Materials and Methods). Additionally, there was an effect of training depth on explicit knowledge (F_1, 54_ = 4.220, p = 0.045, η_p_^2^ = 0.072), indicating that free-recall accuracy was better with increased practice. However, there was no interaction of training depth and explicit cues (F_1, 54_ = 0.315, p = 0.577, η_p_^2^ = 0.006). The explicit cueing manipulation successfully resulted in increased knowledge of explicit sequences, independent of the amount of practice.

**Figure 5.**
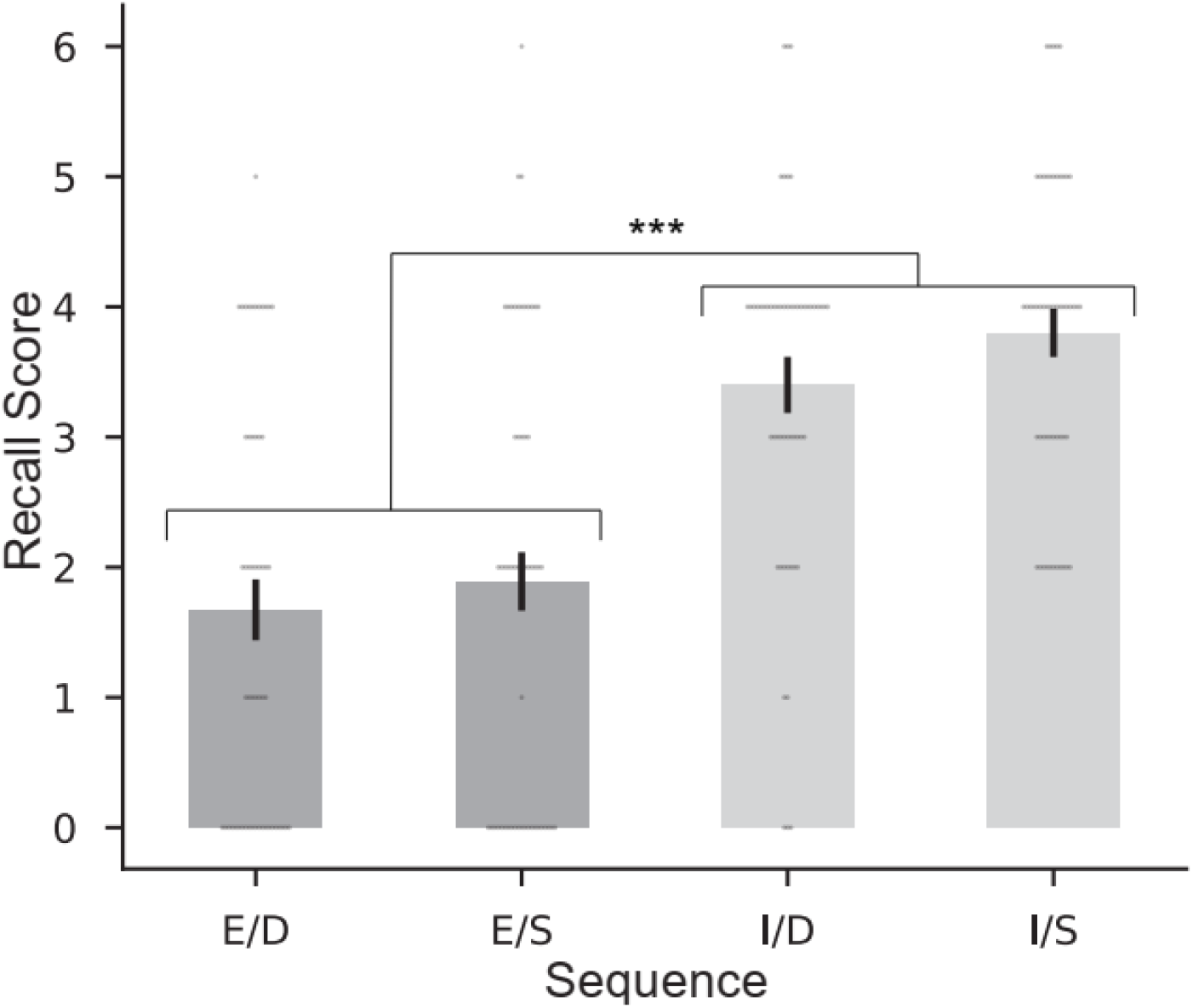
Explicit knowledge of sequences as assessed by free recall. Following the experiment, participants were prompted to type each sequence from memory given either the colored sequence cue (Explicit sequences) or the first item (Implicit sequences). Recall error is defined as the number of transpositions and/or substitutions needed to transform typed sequences into the actual sequences, with zero signifying perfect memory. As expected, explicit knowledge was greater for the explicitly trained sequence. Gray dots indicate individual subjects. Error bars indicate SEM^norm^. (E/D - Deeply trained, Explicit; I/D - Deeply trained, Implicit, E/S - Shallowly trained, Explicit, I/S - Shallowly trained, Implicit; *** = p < 0.001).

#### 2.2.6. Degree of explicit knowledge related to overall performance in motivated context

To assess in more detail how varying amounts of explicit knowledge affects performance, we conducted two separate analyses on the relationship between MT during Reward and participants’ accuracy in free-recall. If sequence knowledge simply enhances motivational vigor, one might expect that knowledge should improve performance regardless of whether a pre-cue allows for advanced motor preparation. In this first analysis, we simply calculated the Spearman’s correlation between MT during Reward and recall error. Explicit knowledge was significantly correlated with MT during Reward in both of the explicitly cued sequences (Explicit/Deep: r_s_ = 0.301, p = 0.025, Explicit/Shallow: r_s_ = 0.573, p < 0.001; Figure 6a). These correlations were not significant in the implicitly trained sequences (Implicit/Deep: r_s_ = 0.169, p = 0.216, Implicit/Shallow: r_s_ = 0.090, p = 0.514; Figure 6b). In a second follow-up analysis, we ran a linear mixed effects model to predict MT using cue, training depth, and recall error as predictors with participants as a random factor. Across all sequences, better explicit recall for each sequence was predictive of faster MT (β = 79.36, SE = 23.66, 95% CI = 32.99-125.72, z = 3.36, p = 0.001). However, we did not find evidence that this relationship was stronger for explicit sequences relative to implicit sequences as the interaction between free recall score and cue type (explicit/implicit) was not significant (β = 39.17, SE = 37.24, 95% CI = −112.16-33.82, z = 0.90, p = 0.29). Nonetheless, we found evidence that explicit knowledge of the sequences was strongly related to performance in the explicitly cued conditions.

**Figure 6.**
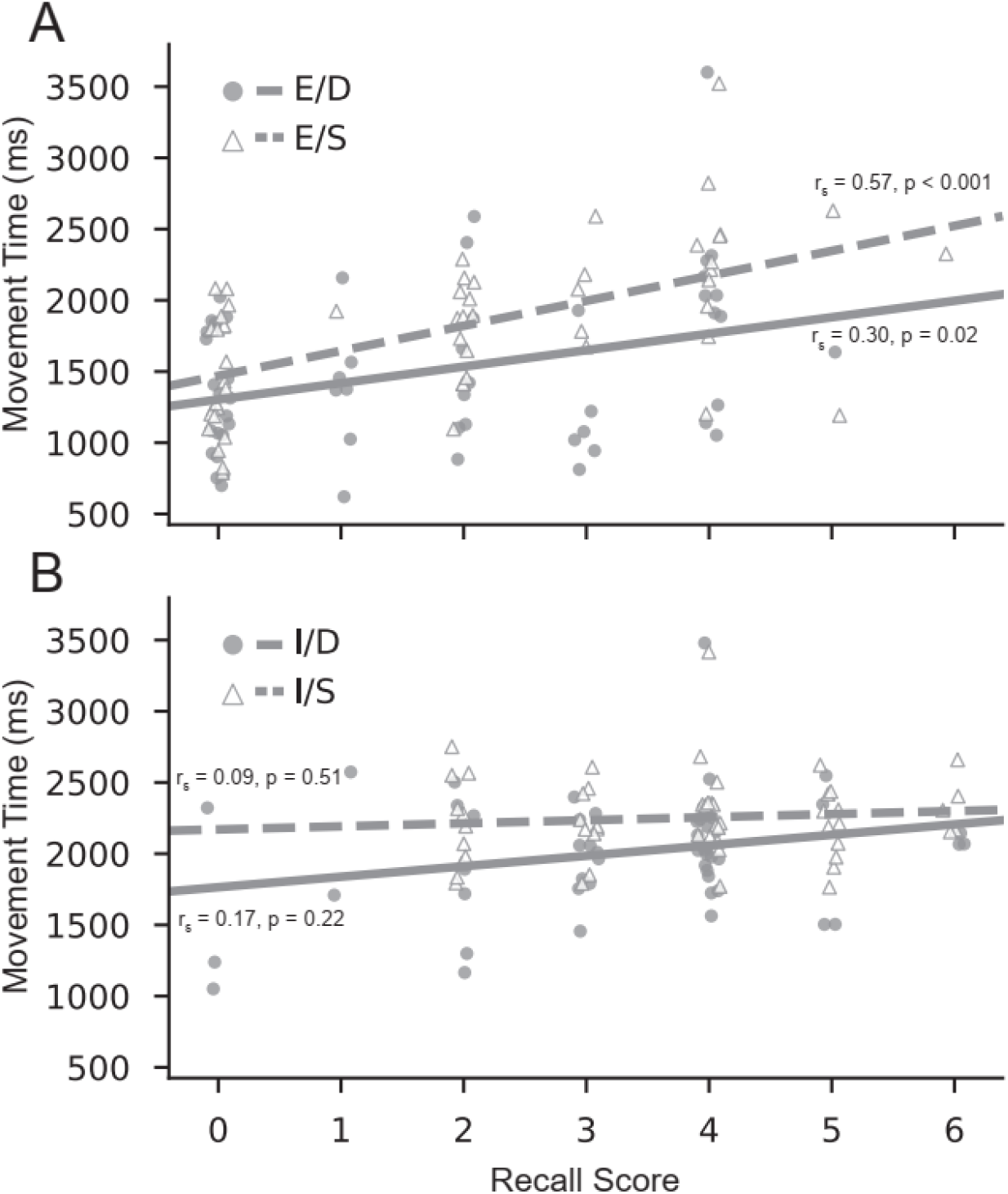
Relationship between explicit knowledge and movement time during the reward phase of the experiment. a) Greater explicit knowledge as assessed by the free recall tests at the end of the experiment was associated with faster movement times during performance with incentives for Explicit sequences. b) There was no significant association between explicit knowledge and performance for Implicit sequences.

#### 2.2.7. Explicit knowledge is predictive of the degree of reward-induced performance enhancement

To further understand the relationship of explicit skill knowledge to the immediate reward-induced performance improvements, we looked at the correlations between explicit knowledge as measured by the free-recall tests and percentage improvement in MT (between T-Last10 and R-First10). We observed a significant correlation between explicit knowledge and MT improvement in both of the Explicit sequences (Explicit/Deep: r_s_ = −0.343, p = 0.010, Explicit/Shallow: r_s_ = −0.332, p = 0.013; Figure 7a) suggesting that greater sequence knowledge allowed for larger incentive-based skill enhancement. We did not observe a significant relationship between sequence knowledge and improvement in MT in either of the implicit sequences: Implicit/Deep (r_s_ = −0.114, p = 0.408), and Implicit/Shallow (r_s_ = −0.217, p = 0.111; Figure 7b). Using a linear mixed effects model with cue (explicit/implicit), training depth, and recall error as predictors and participants as a random factor, we found that greater explicit knowledge was significantly predictive of larger reward-related enhancements in performance (β = 0.013, SE = 0.005, 95% CI = 0.003-0.024, z = 2.50, p = 0.013). However, we did not observe a significant interaction between cue and explicit knowledge (β = −0.003, SE = 0.008, 95% CI = −0.019-0.013, z = 0.378, p = 0.71). These results suggest that explicit sequence knowledge modulates performance improvements due to incentives.

**Figure 7.**
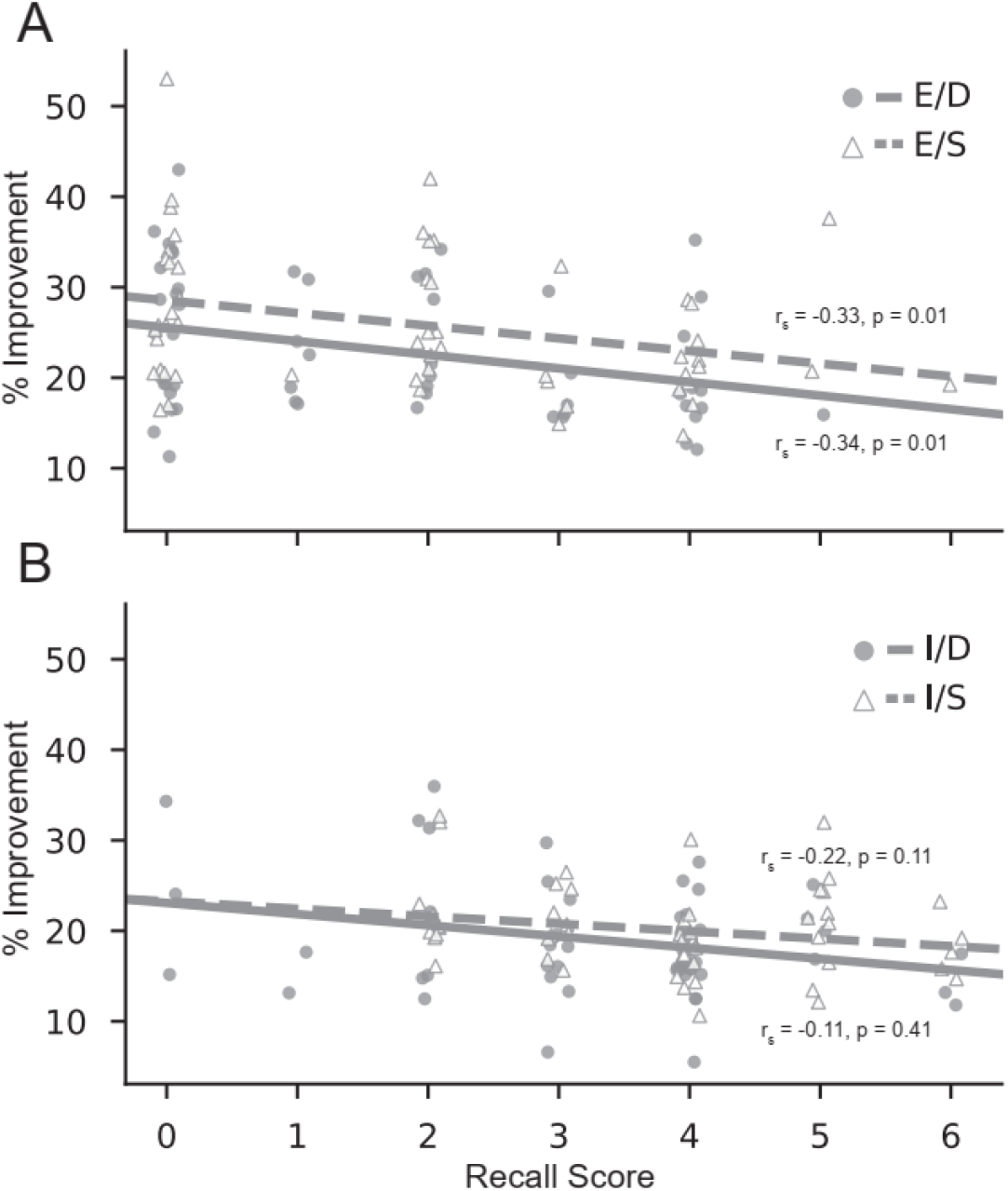
Relationship between explicit knowledge and immediate incentive-based performance improvements. Participants with greater explicit knowledge of the shallowly trained, explicit sequence showed the largest decreases in movement time following the introduction of performance-contingent incentives.

In order to further investigate how explicit knowledge interacts with the ability to plan movements due to explicit cues, we performed an additional analysis looking specifically at the subset of participants (N=17) who had accurate recall of implicit sequences (recall error <2). Despite having explicit knowledge of implicit sequences, these individuals still showed larger enhancement for explicit sequences than implicit sequences (main effect of Cue: F_1,16_ = 6.59, p = 0.021, η_p_^2^ = 0.292). We did not observe a main effect of depth (F_1,16_ = 0.535, p = 0.475, η_p_^2^ = 0.032) nor an interaction (F_1,16_ = 0.132, p = 0.721, η_p_^2^ = 0.008). This suggests that explicit knowledge enables greater incentive-based performance enhancements when explicit cues are available.

#### 2.2.8. Explicit knowledge as measured by recognition tests was not predictive of performance

In addition to the free-recall tests, we assessed knowledge of the sequences after the experiment by assessing how well participants could distinguish between Untrained sequences and the four trained sequences in a recognition test. Participants were able to identify all the trained sequences as “old” in this test (one sample t-tests: all p ≤ 0.01), but there was no main effect of cue nor training depth on recognition score (all p > 0.25). Somewhat surprisingly, accuracy in the free-recall test and recognition test were not correlated (all p’s > 0.15). Additionally, recognition scores were not correlated with MT performance in the Reward phase of the experiment (all p’s > 0.15).

## 3. EXPERIMENT 2

Many of the analyses in Experiment 1 were exploratory. In order to confirm the results from experiment 1, we ran a direct replication that was pre-registered using the Open Science Framework (OSF) (https://osf.io/rs2ha/).

### 3.1. Experiment 2 Materials and Methods

#### 3.1.1. Participants

35 right-handed subjects participated in Experiment 2 (19 females and 16 males, all right-hand dominant, mean age = 21.11, SD = 3.08 years). An additional 6 participants were consented, but then excluded because of ineligibility for the study or experimenter error. One participant was excluded from the recognition questionnaire analysis due to experimental error. All participants were paid $10/hr + performance bonuses for their participation and provided written informed consent.

All methods and procedures were identical to those reported in Experiment 1.

### 3.2. Experiment 2 Results

#### 3.2.1. Explicit cues and increased practice led to better learning and performance in Training

The results from Experiment 2 confirmed the effectiveness of explicit cues in enhancing motor sequence learning relative to implicit learning (Figure 8). Explicit sequence cues again led to faster MT (main effect of cue: F_1,33_ = 116.25, p < 0.001, η_p_^2^ = 0.779), a faster learning rate (cue x block interaction: F_4.26,140.55_ = 5.65, p < 0.001, η_p_^2^ = 0.146), and faster RT at the start of each sequence (main effect of cue: F_1,33_ = 23.65, p < 0.001, η_p_^2^ = 0.417). While there was again no effect of Explicit cues on accuracy rate (main effect of cue: F_1,34_ = 0.696, p = 0.410, η_p_^2^ = 0.020), we did find a main effect of block on accuracy rate (F_7,238_ = 2.80, p = 0.008, η_p_^2^ = 0.076; see Figure S2). This effect was driven by the fact that participants were significantly more accurate during the first block of training when movement times were still relatively slow (block 1 vs block 2: t_34_ = 3.17, p = 0.002). Accuracy rate did not significantly differ between any subsequent consecutive blocks (range of t-values = 0.15-1.32, range of p-values = 0.19-0.88). Explicit cues improved both learning and performance during training.

**Figure 8.**
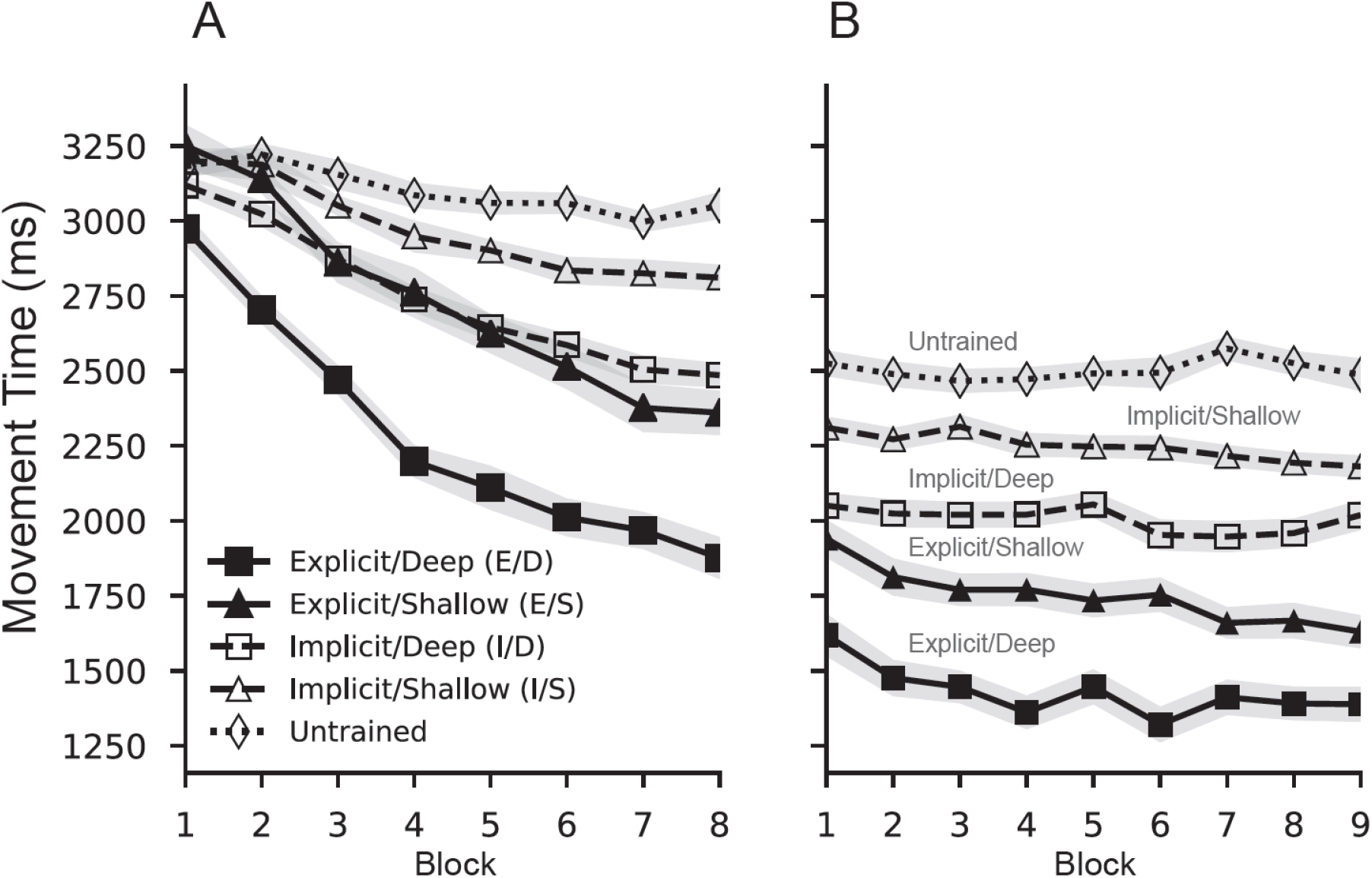
Mean movement time (MT) for correctly executed sequences in Experiment 2. MT was measured as the total time between the first and last item in each sequence. a) Explicit sequence cues and increased training were both associated with faster rates of learning. b) Following the introduction of performance-contingent monetary incentives, movement times immediately decreased for all sequences. Shaded areas indicate SEM^norm^. (E/D - Deeply trained, Explicit; I/D - Deeply trained, Implicit, E/S - Shallowly trained, Explicit, I/S - Shallowly trained, Implicit).

We also validated the effect of practice (depth of training) on performance. MT and RT were both faster for the deeply trained sequences (main effects of depth: F_1,33_ = 31.50, p < 0.001, η_p_^2^ = 0.488, F_1,33_ = 15.26, p = 0.001, η_p_^2^ = 0.316, respectively), learning rate was greater (depth x block interaction: F_2.87, 94.83_ = 15.971, p < 0.001, η_p_^2^ = 0.326), and accuracy rate was higher (F_1,34_ = 72.882, p < 0.001, η_p_^2^ = 0.682). Additionally, we again observed a knowledge type by training depth interaction in MT whereby the benefit of training depth on MT was more pronounced for explicitly cued sequences than for implicitly trained sequences (F_1,33_ = 16.30, p < 0.001, η_p_^2^ = 0.331). We also observed a knowledge type by training depth interaction in accuracy (F_1,34_ = 5.848, p = 0.021, η_p_^2^ = 0.147). The benefit of practice depth on accuracy was greater for implicit sequences. It is possible that increased practice (deep training) contributed to skill by means of faster MT in explicit sequences, whereas in implicit sequences deep training contributed to skill with better accuracy. The results from Experiment 2 replicated the benefit of depth of training on learning and performance.

#### 3.3.2. Immediate reward-related performance enhancement larger for explicitly trained skills

To confirm whether explicit knowledge moderates the effect of motivation on performance through improved motor planning, we again examined the immediate effect of introducing performance-contingent monetary incentives following training. Again, percentage improvement was used in order to normalize enhancement benefits to each individual’s performance at the end of Training. Replicating the results from Experiment 1, participants immediately decreased their MT for all trained sequences and Untrained sequences (see Figure 9; p < 0.001 in one-sample t-tests for all sequences). We again observed a large decrease in MT on Untrained sequences (18%), suggesting that monetary incentives had a global effect on motivational vigor independent from any sequence-specific skill knowledge. Replicating Experiment 1, the performance boost was virtually identical for Untrained sequences and the two implicitly trained sequences (no main effect of sequence identity, explicit sequences excluded: F_1.7,57.9_ = 0.905, p = 0.396, η_p_^2^ = 0.026), while MT improvement for explicitly cued sequences was significantly larger than those for implicitly trained sequences (main effect of cue type: F_1,34_ = 25.607, p < 0.001, η_p_^2^ = 0.43) and the untrained sequence(E/D vs. Untrained: t_34_ = 5.77, p < 0.001; E/S vs. Untrained: t_34_ = 7.00, p < 0.001). None of the above results differed substantially when 5 or 15 trials were used to calculate the immediate effect of incentives (all p’s < 0.001, see Supplementary Results). This again suggests that explicit knowledge in the presence of sequence cues allowed for skill enhancements above and beyond simple increases in overall motor vigor. In Experiment 2, we did not find a main effect of training depth (F_1,34_ = 0.097, p = 0.757, η_p_^2^ = 0.003) nor a depth x cue type interaction (F_1,34_ = 0.038, p = 0.847, η_p_^2^ = 0.001), pointing more strongly to the influence of explicit knowledge and not amount of training. As in Experiment 1, participants performed similarly during the last block of training on the I/D sequence and the E/S sequence (t_34_ = 1.47, p = 0.15), yet the immediate reward-based performance improvement was much larger for the explicitly trained sequence (t_34_ = 3.22, p < 0.005). Additionally, the immediate absolute change in MT in raw ms was larger for E/S than I/D (t_34_ = 3.194, p = 0.003), replicating Experiment 1 and again ruling at that all sequences simply improved by the same raw amount of time. These confirmatory results suggest that incentives improve overall movement speed, but lead to additional enhancements of motor performance in skills with explicit knowledge.

**Figure 9.**
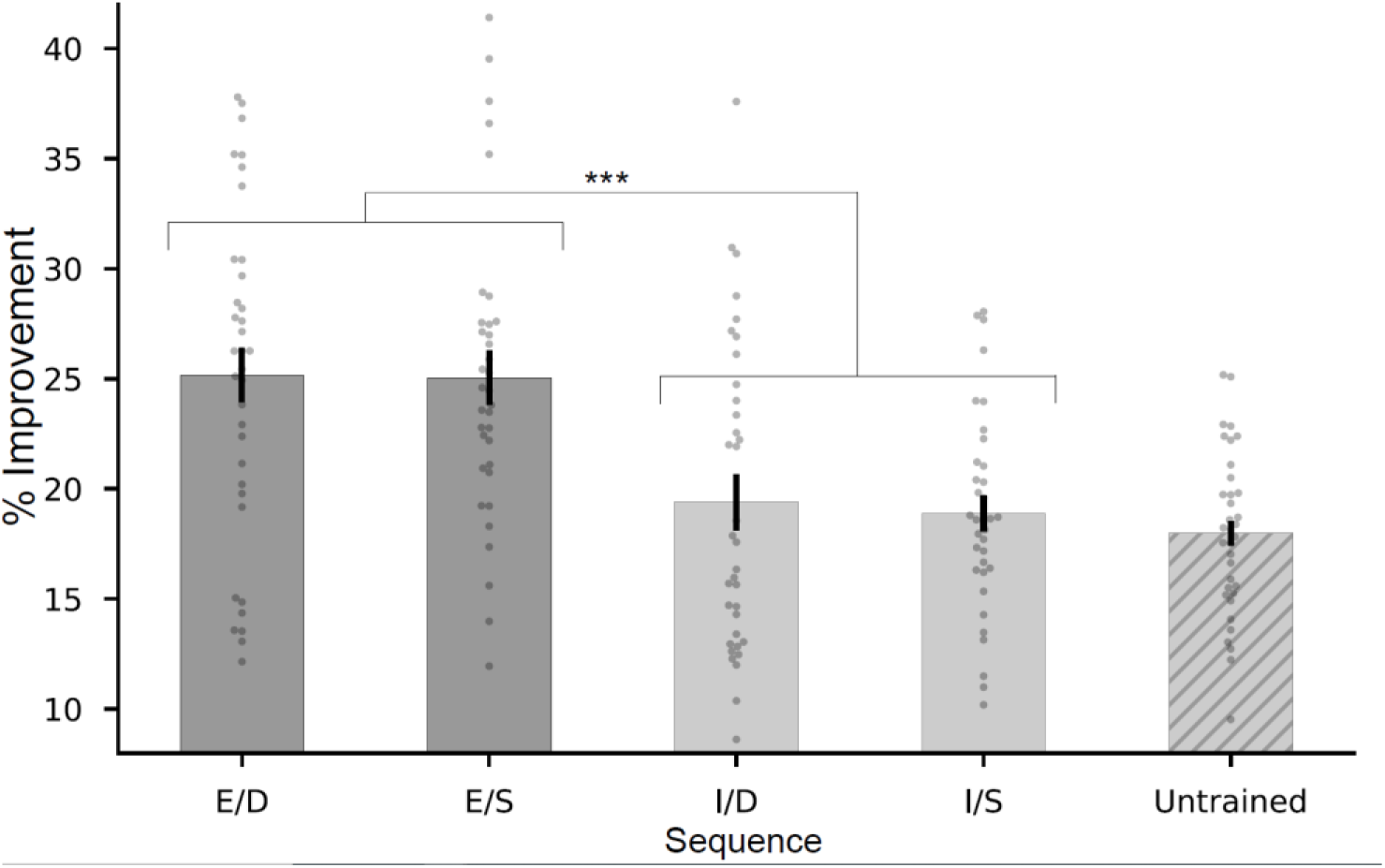
Replication of incentive-based performance improvements. In Experiment 2, performance-contingent monetary incentives led to larger gains in performance on explicitly trained sequences relative to implicit and untrained sequences. Error bars indicate SEM^norm^. (E/D - Deeply trained, Explicit; I/D - Deeply trained, Implicit, E/S - Shallowly trained, Explicit, I/S - Shallowly trained, Implicit).

As in Experiment 1, these effects also cannot be explained by the fact that participants are relatively faster at the end of training for the explicitly cued sequences. We did not find a statistically significant relationship between skill level, as indexed by MT, at the end of training and the size of the reward-related performance enhancement for any of the sequences (E/D: r_s_ = −0.12, p_uncorrected_ = 0.5; E/S: r_s_ = 0.08, p_unc_ = 0.64; I/D: r_s_ = 0.02, p_unc_ = 0.90; I/S: r_s_ = −0.19, p_unc_ = 0.28; Untrained: r_s_ = −0.31, p_unc_ = 0.08; all p_corrected_’s > 0.4 after Holm-Bonferroni correction).

The introduction of monetary incentives again led to faster RT to the first item of each sequence (p < 0.001 in one-sample t-tests for all sequences). We again found no significant effects of cue type or training depth on this RT improvement suggesting that this effect was similar across all trial types (main effect of cue type: F_1,34_ = 1.50, p = 0.229, η_p_^2^ = 0.042; main effect of training depth: F_1,34_ = 0.86, p = 0.361, η_p_^2^ = 0.025; depth x cue interaction: F_1,34_ = 0.19, p = 0.67, η_p_^2^ = 0.005). This suggests that the performance improvement for explicit sequences is in part due to enhancements in movements beyond the first item in the sequence.

We again examined percentage improvement within each IKI between explicit and implicit sequences to assess the enhancements of motor planning. Similar to Experiment 1, we observed more pronounced performance enhancements for earlier items in each sequence (IKIs 2 and 3, both p’s < 0.006, Bonferroni-corrected; see Figure 10), consistent with the idea that the introduction of monetary incentives led to enhanced planning in explicit sequences. We additionally observed larger improvements in IKIs at the end of the sequence (IKIs 7 and 8, both p’s < 0.02, Bonferroni-corrected; all other p’s > 0.6). For the implicitly trained sequences, the incentive-related improvement among all IKIs was similar and not significantly different from the improvement seen for the first item in the sequence (all p’s > 0.05, Bonferroni-corrected). This is again consistent with the conclusion that all items in implicitly trained sequences see a global enhancement in movement vigor at the introduction of monetary incentives.

**Figure 10.**
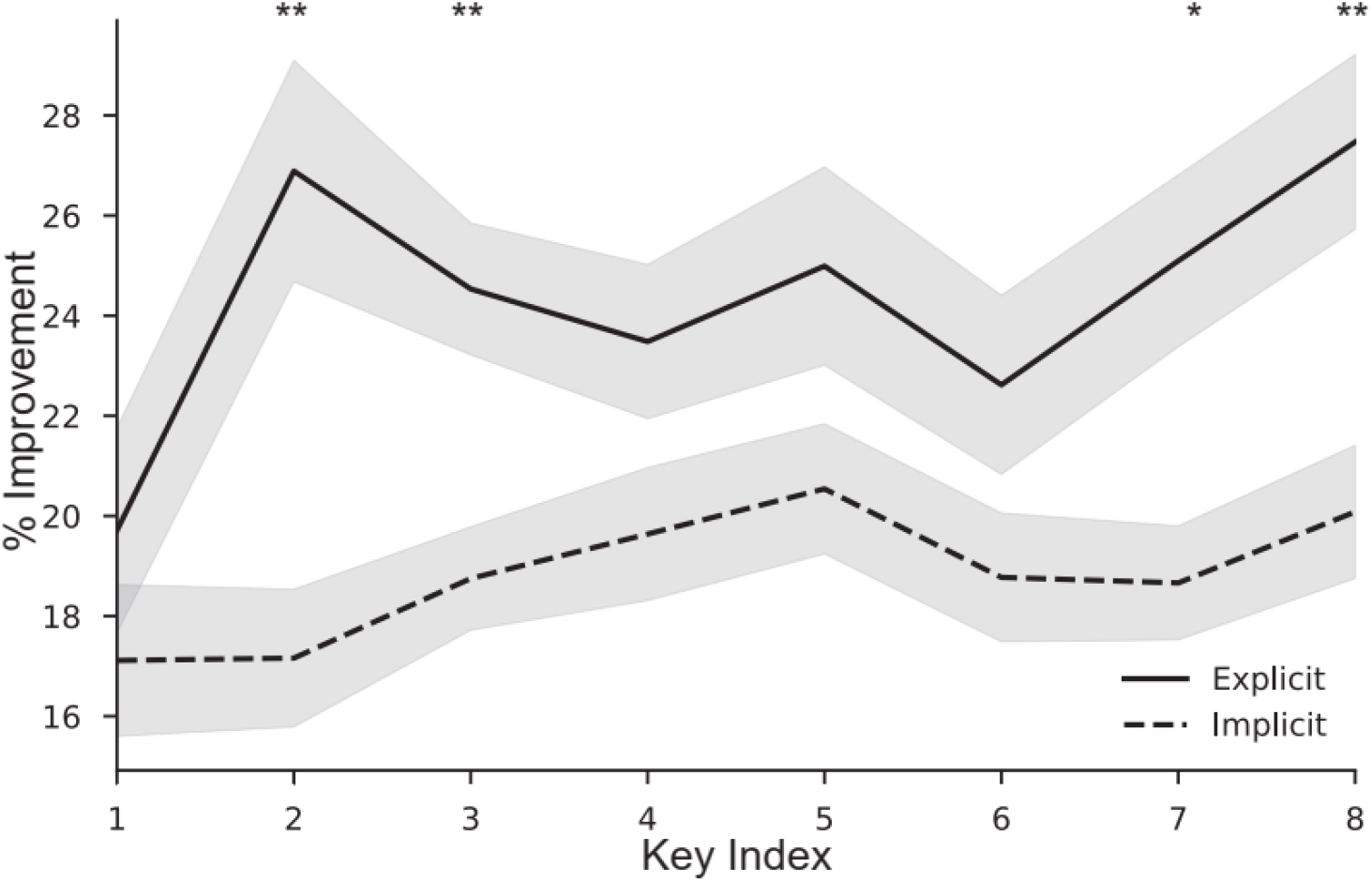
Incentive-based performance improvements for individual items within a sequence in Experiment 2. Although all individual movements within a sequence benefitted from the introduction of performance-contingent monetary incentives, items earlier in the sequence (IKIs 2 and 3) and at the very end of the sequence (IKIs 7 and 8) showed significantly larger boosts for explicitly cued sequences. (** = p < 0.01, * = p < 0.05, Bonferroni-corrected. No other significant differences.)

#### 3.3.3. Better performance in explicitly cued sequences and deeply trained sequences was robust in motivated context

To confirm the performance benefits of explicit knowledge and increased practice persisted in contexts with increased motivation, we again looked at overall performance in the Reward phase. Explicit cues led to faster MT in Reward (F_1,34_ = 59.413, p < 0.001, η_p_^2^ = 0.636). Explicit cues also led to faster RT (F_1,34_ = 70.801, p < 0.001, η_p_^2^ = 0.676). Replicating Experiment 1, the benefit of explicit cues in MT and RT persisted after incentives were introduced. MTs were faster in more heavily practiced sequences during Reward (main effect of training depth: F_1,34_ = 85.915, p < 0.001, η_p_^2^ = 0.716). The same effect was observed in RT (F_1,34_ = 5.932, p = 0.020, η_p_^2^ = 0.149). Replicating Experiment 1, the overall benefits of explicit cues and depth of training observed during Training remained with the addition of monetary incentives.

#### 3.3.4. Accuracy increased with higher incentive values

We next assessed whether performance accuracy in Experiment 2 was sensitive to incentive magnitude in the Reward phase. Participants were somewhat sensitive to incentive magnitude; in contrast to the results seen in Experiment 1, they were more accurate as reward value grew large (main effect of incentive magnitude: F_2,68_ = 3.19, p = 0.047, η_p_^2^ = 0.086). Unlike Experiment 1, we did not observe a reward by training depth interaction (F_2,68_ = 0.234, p = 0.792, η_p_^2^ = 0.007). Given the effect of incentive magnitude on performance was inconsistent and relatively weak across the two experiments, we will refrain from discussing them further.

Perhaps due to the use of strict time limits, shallowly trained sequences were again more accurate during the Reward phase (main effect of training depth: F_1,34_ = 10.46, p < 0.005, η_p_^2^ = 0.235) as were implicit sequences (main effect of cue type: F_1,34_= 7.90, p < 0.01, η_p_^2^ = 0.189). Additionally, there were no significant interactions between reward, training depth, or cue type (all p > 0.15). Neither explicit sequence knowledge nor the depth of training affected the impact of incentive magnitude on performance.

#### 3.3.5. Explicit knowledge greater for explicitly cued sequences

As expected, participants were able to recall explicitly cued sequences more accurately than implicitly trained ones (F_1,34_ = 49.790, p < 0.001, η_p_^2^ = 0.594). Unlike Experiment 1, there was no effect of training depth on explicit knowledge (F_1,34_ = 0.004, p = 0.949, η_p_^2^ = 0.000), indicating that increased practice and exposure did not result in greater explicit sequence knowledge (see Figure S3). Experiment 2 confirms that the explicit cueing manipulation successfully resulted in increased knowledge of explicit sequences, independent of the amount of practice.

#### 3.3.6. Degree of explicit knowledge related to performance in motivated context

To assess in more detail how varying amounts of explicit knowledge affects performance, we again computed correlations between MT during Reward and participants’ accuracy in free-recall. Congruent with Experiment 1, explicit knowledge was significantly correlated with mean MT during Reward in both of the explicitly cued sequences (E/D: r_s_ = 0.586, p < 0.001, E/S: r_s_ = 0.619, p < 0.001). Furthermore, this correlation was not significant in the implicitly trained sequences (I/D: r_s_ = 0.239, p = 0.166, I/S: r_s_ = −0.024, p = 0.891). Explicit knowledge of the sequences was again strongly related to performance in only the explicitly cued conditions. We again ran a linear mixed-effects model to predict MT with cue, depth, and recall as predictors with participants as a random factor. Replicating the results from Experiment 1, greater explicit knowledge was significantly predictive of faster MT during the Reward phase of the experiment (β = 140.78, SE = 27.93, 95% CI = 86.04-195.53, z = 5.04, p < 0.001; Figure S4). We also observed a significant interaction between cue and explicit knowledge such that the relationship between MT and explicit knowledge was stronger for explicitly cued sequences (β = 82.57, SE = 40.13, 95% CI = 3.92-161.23, z = 2.06, p = 0.04).

#### 3.3.7. Explicit knowledge is predictive of the degree of reward-induced performance enhancement

To confirm explicit skill knowledge is related to immediate reward-induced performance improvements, we again examined the relationship between explicit knowledge and the immediate performance gains seen in response to the introduction of monetary incentives. We again observed a significant correlation between explicit knowledge and MT improvement in the Explicit/Shallow sequence (r_s_ = −0.47, p = 0.004) suggesting that greater sequence knowledge allowed for larger incentive-based skill enhancement (See Figure S5). Again, this relationship did not hold for either of the implicit sequences: Implicit/Deep (r_s_ = −0.087, p = 0.62), and Implicit/Shallow (r_s_ = 0.055, p = 0.75). Interestingly, we did not find a significant relationship between sequence knowledge and the level of MT improvement for the Explicit/Deep sequence (r_s_ = −0.028, p = 0.87), however variability across participants was low as many participants (18 total) had perfect knowledge of this sequence (i.e. edit-distance of 0). If these participants with perfect knowledge are excluded from the analysis, we again observe a significant relationship between the level of explicit knowledge and the size of incentive-based performance improvement in MT (r_s_ = −0.55, p = 0.02). These results again suggest that enhancements induced by performance-based incentives are magnified by explicit knowledge in the presence of sequence cues.

Just like in Experiment 1, several participants achieved implicit recall scores of 2 or better (N=15). Replicating Experiment 1, we observed a main effect of cue on MT improvement (F_1,14_ = 4.187, p = 0.060, η_p_^2^ = 0.230) in this group of subjects. We did not observe a main effect of training depth (F_1,14_ = 0.554, p = 0.469, η_p_^2^ = 0.038) nor an interaction (F_1,14_ = 1.924, p = 0.187, η_p_^2^ = 0.121). This again suggests explicit knowledge contributed to greater performance enhancements for the explicitly cued sequences.

#### 3.3.8. Recognition tests are weak predictors of performance

Just as in Experiment 1, participants were able to identify all the trained sequences as “old” in this test (one sample t-tests: all p ≤ 0.01), and there was no main effect of cue nor training depth on recognition score (all p > 0.5). Accuracy in the free-recall test and recognition test was correlated in the Explicit/Shallow sequence (r_s_ = −0.50, p = 0.0029), but not in any other sequence (all p > 0.5). Across the two experiments, there does not seem to be much evidence that the ability to recognize learned sequences is related to strong free recall of the sequences.

Unlike in Experiment 1, MT and recognition score were correlated in two of the sequences: Explicit/Shallow (r_s_ = −0.4, p = 0.018) and Implicit/Deep (r_s_ = −0.41, p = 0.017). Correlation between Reward MT and recognition score was not significant in Explicit/Deep (r_s_ = −0.084, p = 0.64) nor in Implicit/Shallow (r_s_ = −0.3, p = 0.084). Given the variability in the relationship between motivated performance and explicit knowledge as assessed by the recognition tests across the two experiments, we refrain from making definitive claims regarding these results.

## 4. Discussion

We sought to investigate whether monetary incentives enhance skilled motor performance by simply increasing the motivational vigor of movements, improving movement planning, or both. In our task, participants learned to perform four separate sequences, two of which were explicitly cued to allow for advanced planning. Explicit cues led to a faster rate of skill acquisition and greater levels of skill knowledge at the end of the experiment. Immediately after the introduction of performance-contingent monetary incentives, we observed a global boost in performance in the execution of all trained and Untrained (random) sequences. However, the magnitude of incentive-based performance enhancement was much larger for the explicitly cued sequences, which allow for the pre-planning of movements. Furthermore, the size of this performance boost on explicitly cued sequences increased with the level of explicit knowledge attained by our participants. Additionally, we observed the greatest improvement in movements toward the beginning and at the end of each sequence suggesting more efficient chunking of explicit sequences. These effects were all replicated in a separate, pre-registered experiment. These results provide evidence that performance-contingent monetary incentives lead to improvements in both motor execution and motor planning in motor sequencing skills.

### 4.1. Reward leads to improvements in action execution andselection

There are numerous studies that show that training improves the execution of trained motor sequences (see Krakauer, Hadjiosif, Xu, Wong, & Haith, 2019 for a comprehensive review). However, training in sequence learning tasks, such as the DSP task used here, have also been shown to improve the execution of novel sequences or randomly generated sequences (e.g. Nissen & Bullemer, 1987). This sequence-independent improvement likely relies on improvements in motor execution and simple action selection. That is, participants both move more quickly and improve their ability to map visual stimuli to the correct motor response (i.e. finger press). In our task, the introduction of performance-contingent monetary incentives clearly led to enhancements in motivational vigor that improved one or both of these processes. Participants in both Experiment 1 and Experiment 2 were immediately able to speed up their responses to Untrained sequences by almost 20%. Performance on the implicitly trained sequences improved to a similar extent. Furthermore, we observed the same level of incentive-related enhancement for all individual elements of implicitly trained sequences. It is likely that improvements on implicit sequences were likewise driven by enhancements in either simple execution, action selection in response to each item in the sequence, or both.

This finding of incentive-induced improvements in the execution of trained and novel motor sequences adds to the significant literature demontrating that the availability of reward results in faster, more accurate movements (Manohar et al., 2015; Manohar, Finzi, Drew, & Husain, 2017; Summerside et al., 2018). Prior work has shown that rewards can speed up both simple reaction time and action selection with a simple stimulus-response mapping (Mir et al., 2011; Ramnani & Miall, 2003). We cannot disentangle which of these two processes is more affected by motivation in our paradigm. Although several studies have shown that the introduction of performance-contingent rewards also benefits the execution of motor skills (Mosberger et al., 2016; Wachter et al., 2009), the current results suggest that for implicitly trained skills improvement is simply the result of a global increase in movement speed and not skill-specific.

### 4.2. Potential improvements in planning for explicitly trained sequences

The positive effect of prospective rewards on motor execution and action selection almost certainly contributed to the enhanced performance seen for explicitly cued sequences as well. However, the size of the immediate reward-related performance enhancement was larger for explicitly cued sequences than implicitly trained and Untrained sequences. Specifically, this additional enhancement occurred for individual items early on in each sequence and at the very end of each sequence. In sequence learning experiments, most analyses of how individual items are chunked together assume that the second item and the last item in each sequence cannot be the start of a new motor chunk (Bo & Seidler, 2009; Kennerley, Sakai, & Rushworth, 2004). Though the large differences in movement time between conditions made us unable to employ more traditional chunking analyses on our data, it appears that the greatest reward-related enhancement occurred for items *within* a motor chunk rather than the start of chunk. These results are consistent with the hypothesis that the availability of reward and increased motivation also conferred a benefit to the movement planning afforded by the combination of explicit knowledge and the sequence cues.

It is important to note that although initial RT to the first item in each sequence was faster overall for explicitly trained sequences, we did not see any difference between Implicit and Explicit skills in the size of the reward-related boost for initial RT. If planning were enhanced by rewards and additional items could be planned in advance, it seems plausible that this would be reflected in initial RT as well. However, this is complicated by the fact that planning to produce multiple items in advance can actually slow RT to the first item (e.g. Henry & Rogers, 1960) and longer planning times can enhance the speed of subsequent movements (Ariani & Diedrichsen, 2019). It’s possible that the effects of reward enhancement and increased planning on RT essentially canceled each other out with regard to any improvement in RT measured here.

Many studies point to the importance of cognitive control processes in explicit sequence learning (Ashe, Lungu, Basford, & Lu, 2006; Bo & Seidler, 2009; Seidler, Bo, & Anguera, 2012; Unsworth & Engle, 2005). Specifically, larger working memory capacity and proactive control strategies are known to be predictive of the level of skill improvement during training. It is thought that the role of these processes in sequence learning is to aid in pre-planning movements and to aid in the formation of motor chunks. In cognitive tasks, the prospect of reward has been recently shown to lead to enhancements in these same cognitive control processes (Botvinick & Braver, 2014; Jimura et al., 2010; Krawczyk, Gazzaley, & D’Esposito, 2007; Padmala & Pessoa, 2011). It stands to reason that explicit sequence performance in our task benefitted both from reward-related enhancement in cognitive control processes as well as simple increases in motivational vigor. Although several studies have shown that movement planning in the context of a single button press can be enhanced by the availability of reward (Mir et al., 2011; Ramnani & Miall, 2003), here we show this same enhancement can also improve the execution of a more complex skill.

The level of explicit knowledge our participants had about a particular explicitly cued sequence was correlated with the magnitude of reward-related enhancement in performance. Similarly, average movement time throughout the reward phase of the experiment was also correlated with explicit knowledge. Although some participants clearly gained at least some explicit knowledge of the implicitly trained sequences, we did not observe strong evidence of the same knowledge-performance links for these sequences. It is possible that the benefit of explicit knowledge could only be fully realized when sequence cues were available. While recent work suggests that improvements in performance on implicit sequencing skills can be attributed to improvements in movement planning (Ariani & Diedrichsen, 2019), it does not seem that reward amplified online movement planning for implicit skills in our study. We observed similar improvement for all items within each implicitly trained sequence, and this level of improvement was indistinguishable from the improvement seen for Untrained for which online planning was impossible. This again suggests that the benefit of skill knowledge on performance is amplified by reward, but only when the explicit planning of movements is possible. In the context of simple movements, there is some prior work that also suggests that the speed of motor execution is enhanced by reward specifically when the movement can be planned in advance (Mir et al., 2011; Ramnani & Miall, 2003). Although we show here that even unplanned movements (implicit and untrained sequences) can be executed more quickly when rewards are available, there is clearly an added benefit to movement planning.

### 4.3. Knowledge: increased motivation or planning?

Some researchers have suggested that the benefit produced by explicit knowledge in motor sequencing tasks is not related to motor planning, but rather that knowledge acts to globally increase motivational vigor (Wong, Lindquist, Haith, & Krakauer, 2015). In other words, knowledge functionally acts as a reward cue, leading to faster and more accurate movements. This hypothesis can indeed explain why participants in our experiments are faster in performing explicitly cued sequences relative to implicit sequences at the end of training. Along this line of thinking, implicit sequences are performed more slowly simply because participants lack motivation. The pre-cue and explicit knowledge of the to-be-performed sequence increases motivational vigor and results in faster execution of each individual key-press. However, we do not believe that this account provides a parsimonious explanation of the results seen here. We would expect that the introduction of performance-contingent rewards would have a similar or *larger* immediate effect on the performance of implicitly trained sequences when compared to explicit sequences. That is, if the reason that implicitly trained sequences were performed more slowly toward the end of training was because of a relative lack of motivation, one might expect that an immediate increase in motivation due to the prospect of reward would preferentially benefit implicit skills. Against this hypothesis, we instead observed that explicitly trained sequences received the greatest benefit from the introduction of performance-contingent monetary incentives.

If explicit knowledge does not enable movement planning and instead simply enhances motivational vigor, our results would suggest that the addition of extrinsic rewards produces a multiplicative effect on the enhanced intrinsic motivation induced by explicit knowledge. We find this account implausible for several reasons. First, the dominant account of the interaction between intrinsic and extrinsic motivation suggests that extrinsic rewards actually reduce intrinsic motivation rather than amplifying it (Deci, Richard, & Ryan, 1999). Although there is some work that suggests that intrinsic and extrinsic motivation may be additive (Hendijani, Bischak, Arvai, & Dugar, 2016), this work suggests that the two don’t interact and are sub-additive. Second, we would also expect that the level of explicit knowledge for implicit sequences would similarly be associated with the magnitude of reward-related enhancement. We did not find any evidence for such a relationship. Instead, knowledge was only predictive of an immediate reward-related enhancement when participants received a sequence cue that allowed them to plan movements in advance of execution. Third, it is not the case that motivation induced by both incentives and explicit knowledge served to speed of execution of all movements by a constant amount. We observed larger reward-related enhancements in raw movement time in an explicit sequence and implicit sequence which were previously matched in performance at the end of training. Finally, our analysis of individual key-press RTs suggests that the additional enhancement seen in explicitly trained sequences occurs for items toward the beginning and at the end of each sequence. If explicit knowledge itself globally enhances motivational vigor, it is unclear why we would observe differences in reward-related enhancement among successive key-presses within a sequence. With that said, future work using an independent measure of movement planning would likely be fruitful..

### 4.4. Immediate incentive-dependent enhancement is unlikely to be due to rapid consolidation

For all participants in our studies, there was a short break (<5 min) between the end of training and the introduction of monetary incentives while they were given instructions regarding the reward phase of the experiment. One recent study has suggested that it is possible performance improvements on sequence learning tasks can occur offline during short breaks via rapid forms of consolidation (Bönstrup et al., 2019). It is possible that the differences in incentive-based performance improvements that we see between implicitly and explicitly trained sequences are due to differences in rapid consolidation rather than the introduction of incentives. While we cannot completely rule out this possibility, we believe this is unlikely for several reasons. First, the study reporting rapid consolidation found that offline performance improvements occurred primarily at the start of training (first ~10 trials) and diminished in size as training progressed. Monetary incentives were introduced much later in our experiment after many trials of training when the size of any offline consolidation benefits should be negligible. Second, while we observe larger improvements for explicitly trained sequences, most studies of skill consolidation have observed smaller offline consolidation benefits for explicit sequences relative to implicit sequences (Robertson, Pascual-Leone, & Press, 2004). Finally, the immediate performance improvement after the introduction of incentives (>20%) that we observed is much larger than the consolidation benefits reported in prior work on offline consolidation over periods of wakefulness (e.g. Bönstrup et al., 2019; Brown & Robertson, 2007), suggesting a separate mechanism.

### 4.5. Limitations in measuring sequence knowledge

In our experiment, explicit knowledge of the sequences was measured with both a free-recall test and a recognition test. In the free-recall test it was impossible to uniquely test explicit knowledge of implicitly trained sequences without giving away at least the first item of the sequence (see Methods). It is therefore possible that the implicit sequences were recalled in a different way than explicit sequences, which were prompted with the associated color cue. However, the edit-distance measure used to assess performance on this test likely overestimates true explicit knowledge of implicit sequences in this scenario.

Interestingly, we did not find any difference in recognition performance between implicitly and explicitly trained skills. This suggests that participants had at least some fragmentary knowledge of implicitly trained sequences that allowed them to distinguish them from untrained sequences. However, skill knowledge as assessed by these recognition tests was not consistently predictive of performance across the two experiments in contrast with the free recall test. Other researchers have noted that recognition performance can be driven by familiarity in the absence of conscious recollection (Destrebecqz & Cleeremans, 2001). The free recall test was perhaps more “process pure” in measuring explicit knowledge of the sequence.

### 4.6. Inconsistent effects of reward magnitude on performance

Although we provide some evidence that participants in our study are somewhat sensitive to the incentive magnitude during the reward phase of the experiment, these effects were inconsistent across the two experiments. In Experiment 1, we observed that performance on shallowly trained sequences followed an inverted-U shape reminiscent of the effects of choking under pressure we have reported previously (Lee et al., 2019; Lee & Grafton, 2015). In Experiment 2 however, we were unable to replicate this finding. Instead, we simply found that accuracy on all sequences improved with increasing incentive size. There are many individual differences that affect how people respond to monetary incentives, which may have led to the mixed results seen here (Chib, De Martino, Shimojo, & O’Doherty, 2012; Lee & Grafton, 2015; Masters, Polman, & Hammond, 1993). Additionally, while the individually set time-limits in our experiments were intended to match difficulty across the different sequences, they may have led to evaluations of each sequence at different points along each participant’s speed-accuracy curve. That is, participants may have been able to prioritize accuracy over speed for certain sequences because of more relaxed time limits. Furthermore, we rewarded subjects if they were successful on a randomly selected trial during the second phase of the experiment. It is possible that this reward scheme may have led to some discounting of the reward magnitude that resulted in additional variability across participants. These factors could have prevented accurate comparisons of the effect of reward across sequences. Given the inconsistency in results across the two experiments, it is difficult to draw firm conclusions on how incentive magnitude affects performance on this task.

### 4.7. Conclusion

Across two experiments employing a sequence learning task paired with performance-contingent incentives, this study showed motivation improves two separate aspects of skilled motor performance. As reported in previous work, incentives invigorate movement execution leading to some performance improvements independent of both skill knowledge and the level of skill obtained. However, increased motivation also serves to enhance the contribution of explicit knowledge to the performance of motor sequencing skills, perhaps via improved motor planning. This finding helps clarify the mechanism by which enhanced motivation improves skilled performance and also highlights the contribution of explicit training and skill knowledge in sequence learning.

